# TRIM5α restriction of HIV-1-N74D viruses in lymphocytes is caused by a loss of cyclophilin A protection

**DOI:** 10.1101/2020.09.29.318121

**Authors:** Anastasia Selyutina, Lacy M. Simons, Angel Bulnes-Ramos, Judd F. Hultquist, Felipe Diaz-Griffero

## Abstract

The core of HIV-1 viruses bearing the capsid change N74D (HIV-1-N74D) do not bind the human protein cleavage and polyadenylation specificity factor subunit 6 (CPSF6). In addition, HIV-1-N74D viruses have altered patterns of integration site preference in human cell lines. In primary human CD4+ T cells, HIV-1-N74D viruses exhibit infectivity defects when compared to wild type. The reason for this loss of infectivity in primary cells is unknown. We first investigated whether loss of CPSF6 binding accounts for the loss of infectivity. Depletion of CPSF6 in human CD4^+^ T cells did not affect the early stages of wild-type HIV-1 replication, suggesting that defective infectivity in the case of HIV-1-N74D is not due to the loss of CPSF6 binding. Based on our previous result that cyclophilin A (Cyp A) protected HIV-1 from human tripartite motif-containing protein 5α (TRIM5α_hu_) restriction in CD4^+^ T cells, we tested whether TRIM5α_hu_ was involved in the decreased infectivity observed for HIV-1-N74D. Depletion of TRIM5α_hu_ in CD4^+^ T cells rescued the infectivity of HIV-1-N74D, suggesting that HIV-1-N74D cores interacted with TRIM5α_hu_. Accordingly, TRIM5α_hu_ binding to HIV-1-N74D cores was increased compared with that of wild-type cores, and consistently, HIV-1-N74D cores lost their ability to bind Cyp A. In conclusion, we showed that the decreased infectivity of HIV-1-N74D in CD4^+^ T cells is due to a loss of Cyp A protection from TRIM5α_hu_ restriction activity.

## INTRODUCTION

Human cleavage and polyadenylation specificity factor subunit 6 (CPSF6) is a nuclear protein that belongs to the serine/arginine-rich protein family. Expression of a cytosolic fragment of CPSF6 [CPSF6(1–358)] was found to potently block human immunodeficiency virus-1 (HIV-1) infection before the formation of 2-long terminal repeat circles [1], and this inhibition of HIV-1 infection correlated with the ability of CPSF6(1-358) to bind to the capsid and prevent uncoating [2-4]. The serial passaging of HIV-1 in human cells overexpressing CPSF6(1-358) resulted in the generation of escape-mutant viruses bearing the N74D capsid change(HIV-1-N74D) [1], and binding studies of HIV-1 capsids with N74D mutations to CPSF6(1-358) demonstrated a lack of binding as the mechanism for escape [1, 2]. Although the overexpression of full-length CPSF6 remained nuclear and did not block HIV-1 infection, these experiments functionally linked CPSF6 to the HIV-1 capsid. Knockdown or knockout of human CPSF6 expression in different human cell lines did change HIV-1 integration site selection [2, 5-8]. Several reports have also suggested that full-length CPSF6 may facilitate the entry of the virus core into the nucleus [8-10]. The lack of a correlation between the loss of CPSF6 binding to HIV-1 and decreased infectivity in human cell lines indicates that cell-type specific differences in the pathways surrounding early replication contribute to these discrepancies, suggesting the need to work in primary human cell models.

This work used human primary cells to examine the role of CPSF6 in HIV-1 replication. Our experiments demonstrated that the HIV-1 capsid mutations N74D or A77V affected the capsid’s ability to interact with CPSF6, as displayed by infectivity phenotypes in human primary peripheral blood mononuclear cells (PBMCs) and CD4^+^ T cells. When compared with the wild-type virus, HIV-1-N74D demonstrated decreased infectivity, but HIV-1-A77V infectivity was less affected.

These different mutant virus infectivity phenotypes suggested that the reduced primary cell infectivity observed in the case of HIV-1-N74D was not likely to be due to a CPSF6-binding defect. To test this hypothesis directly, we challenged CPSF6-depleted human primary CD4^+^ T cells with HIV-1-N74D and HIV-1-A77V. Remarkably, the reduced HIV-1-N74D infectivity in human primary cells did not change with depleted CPSF6 expression, suggesting that the loss of capsid-CPSF6 interactions did not account for the decreased infectivity of this mutant virus. One possibility is that a different protein may be responsible for reduced HIV-1-N74D infectivity. Recently, we and others have demonstrated that cyclophilin A (Cyp A) protects the HIV-1 core from restriction by the human tripartite motif-containing protein 5α (TRIM5α_hu_) in primary CD4^+^ T cells. To test whether TRIM5α_hu_ is involved in decreased HIV-1-N74D infectivity, we challenged TRIM5α_hu_-depleted human primary CD4^+^ T cells with HIV-1-N74D. Interestingly, we observed that TRIM5α_hu_ depletion rescued HIV-1-N74D infectivity, suggesting that TRIM5α_hu_ is responsible for the restriction observed in human primary T cells. Because Cyp A protects the core from restriction by TRIM5α_hu_, we also tested the ability of the N74D mutation-containing capsids to bind to Cyp A. We found that these capsids lost their ability to interact with Cyp A, which may explain the reason that HIV-1-N74D is restricted by TRIM5α_hu_. Overall, our results show that the HIV-1-N74D mutant virus is restricted by TRIM5α_hu_ due to an inability to bind Cyp A.

## RESULTS

### HIV-1-N74D exhibits defective infectivity of primary CD4^+^ T cells

To test the role of capsid-CPSF6 interactions during the infection process, we challenged dog and human cell lines with HIV-1 viruses containing the capsid mutations N74D and A77V (both of which prevent capsid interactions with CPSF6). Infectivity of wild-type and mutant HIV-1_NL4-3_Δenv, pseudotyped with vesicular stomatitis virus G (VSV-G) envelopes expressing green fluorescent protein (GFP) as an infection reporter, were normalized using p24 levels. HIV-1-N74D-GFP viruses showed a defect on infectivity when compared to wild type viruses when infecting the lung human cell line A549 or the Jurkat T cell line (Figure 1). HIV-1-A77V-GFP viruses showed a lesser defect when compared to HIV-1-N74D-GFP viruses. By contrast, the infectivity defect of HIV-1-N74D-GFP viruses was not observed in the canine cell Cf2Th, which do not express a TRIM5α orthologues (Figure 1).

**Figure 1.**
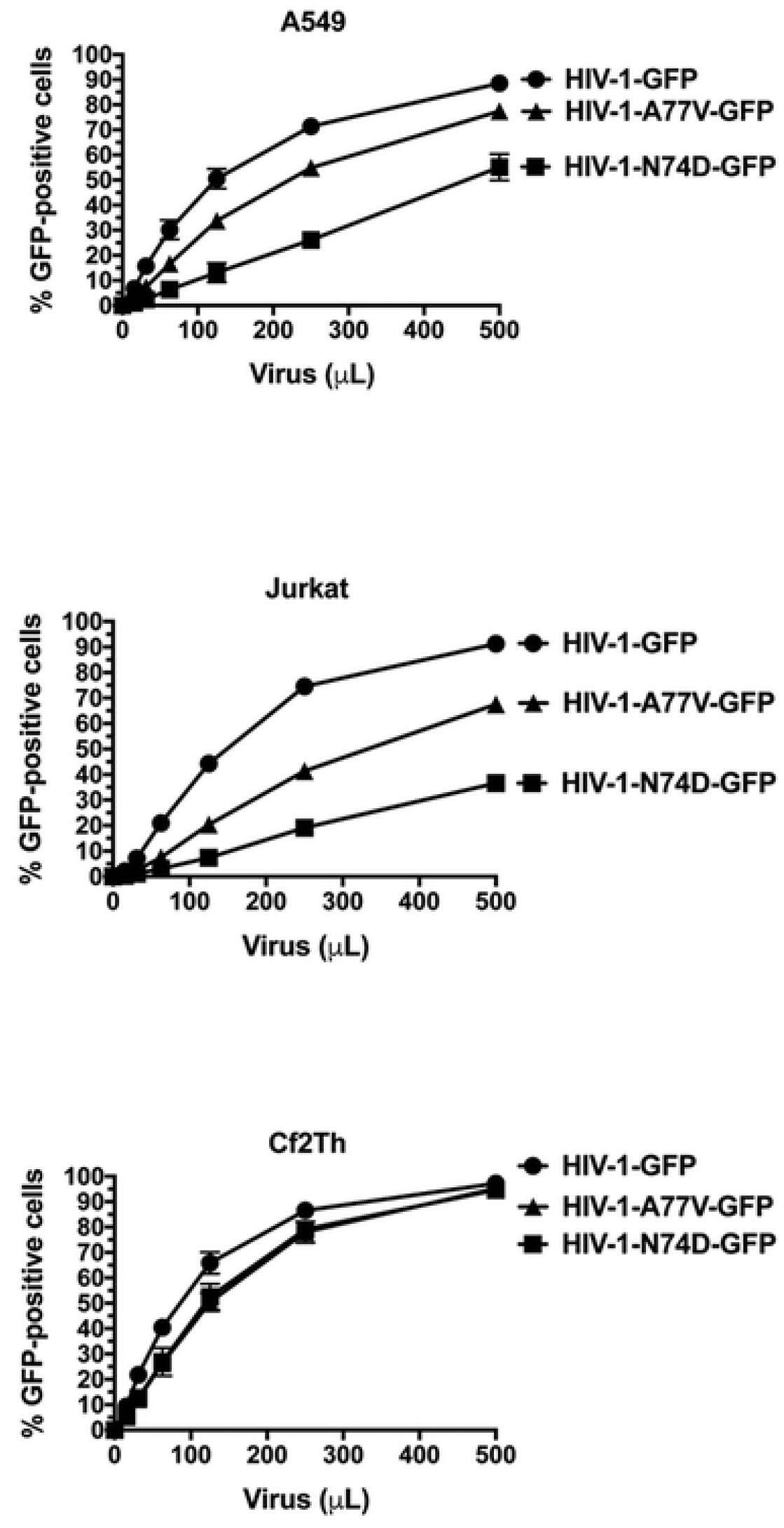
HIV-1-N74D exhibits an infectivity defect in human cell lines but not in dog cell lines. Human lung A549 cells, human Jurkat T cells, or dog thymus Cf2Th cells were challenged with increasing amounts of the indicated p24-normalized WT and mutant HIV-1 viruses. Infectivity was determined at 48 h post-challenge by measuring the percentage of GFP-positive cells. Experiments were repeated three times and a representative experiment is shown.

Next we tested whether these infectivity defects are present in human primary cells. As shown in Figure 2A, HIV-1-N74D-GFP showed a defect in PBMC infections compared with wild-type HIV-1 in at least 3 donors. However, HIV-1-A77V-GFP exhibited a minor infectivity defect when compared with HIV-1-N74D-GFP viruses. Similar results were observed when we challenged human primary CD4^+^ T cells obtained from three independent donors with HIV-1-N74D-GFP and HIV-1-A77V-GFP (Figure 2B). As both N74D and A77V capsid mutants lost their ability to bind to CPSF6 (data not shown), the results suggested that the decreased infectivity of HIV-1-N74D-GFP in primary CD4^+^ T cells is likely due to reasons other than loss of binding to CPSF6.

**Figure 2.**
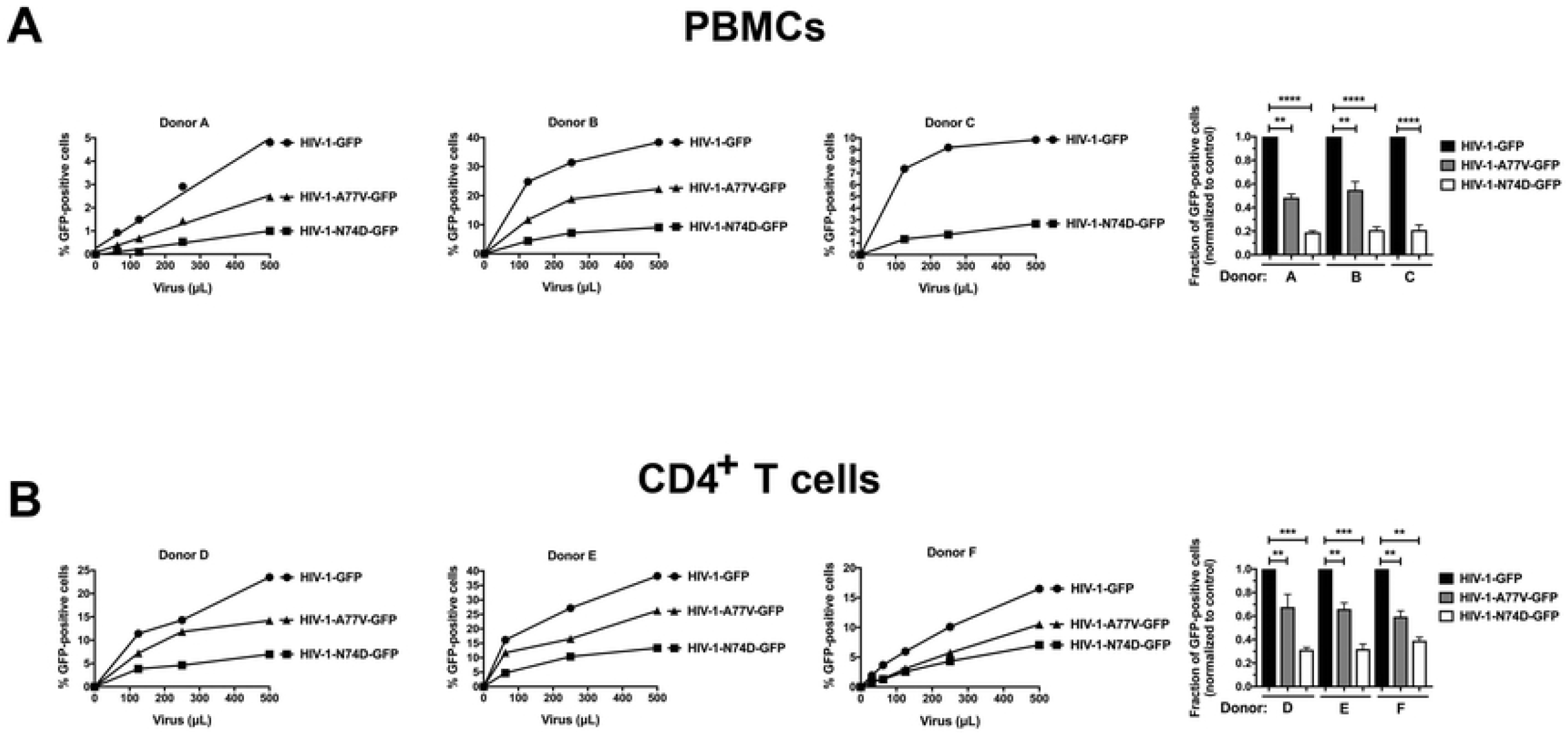
HIV-1-N74D exhibits an infectivity defect in primary PBMCs and CD4^+^ T cells. Human primary PBMCs **(A)** or purified CD4^+^ T cells **(B)** from healthy donors were challenged with increasing amounts of p24-normalized HIV-1-GFP, HIV-1-N74D-GFP, or HIV-1-A77V-GFP. Infectivity was determined at 72 h post-challenge by measuring the percentage of GFP-positive cells. Experiments were repeated three times per donor, and a representative experimental result is shown. Statistical analysis was performed using an intermediate value taken from the infection curves (right panel). ** indicates P-value < 0.001, *** indicates P-value < 0.0005, **** indicates P-value < 0.0001 as determined by using the unpaired t-test.

### Depletion of CPSF6 in human primary CD4^+^ T cells does not affect HIV-1 infectivity

To test the role of CPSF6 in HIV-1 infection of human primary cells, we challenged CPSF6-depleted CD4^+^ T cells with wild-type and mutant HIV-1. As shown in Figure 3A, CRISPR-Cas9 ribonucleoprotein complexes (crRNPs) containing the anti-CPSF6 guide RNA (gRNA) #5 and #6 completely depleted the expression of CPSF6 in human primary CD4^+^ T cells. As a control, we also knocked out the expression of CXCR4. Similar to the results above, CPSF6 depletion did not affect wild-type HIV-1 infectivity in human primary cells (Figure 3B). In addition, depletion of CPSF6 did not affect the infectivity of either HIV-1-N74D-GFP or HIV-1-A77V-GFP. These experiments demonstrated that depletion of CPSF6 in human primary cells did not affect HIV-1 infectivity, suggesting that the reduced infectivity of HIV-1-N74D was not due to blocked virus interactions with CPSF6.

**Figure 3.**
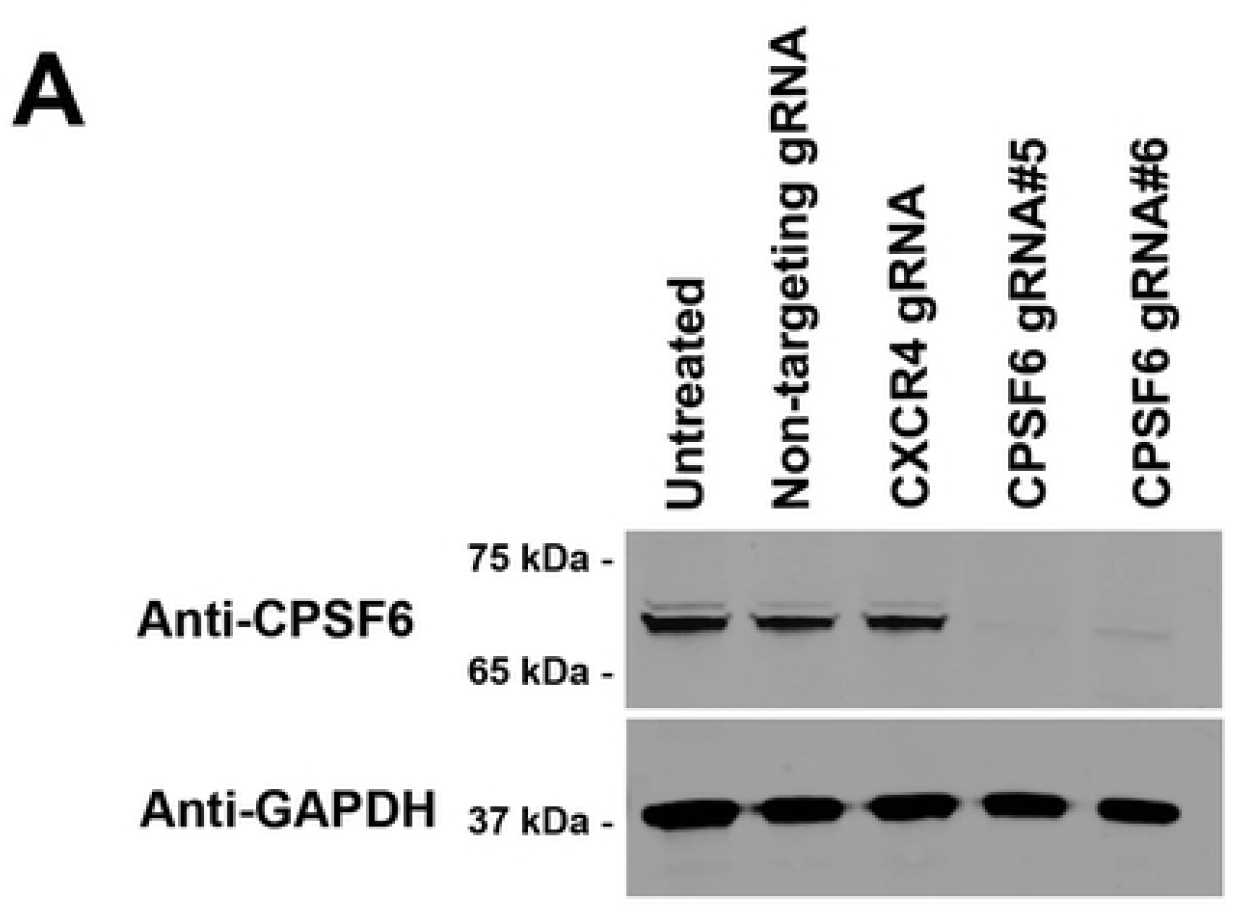
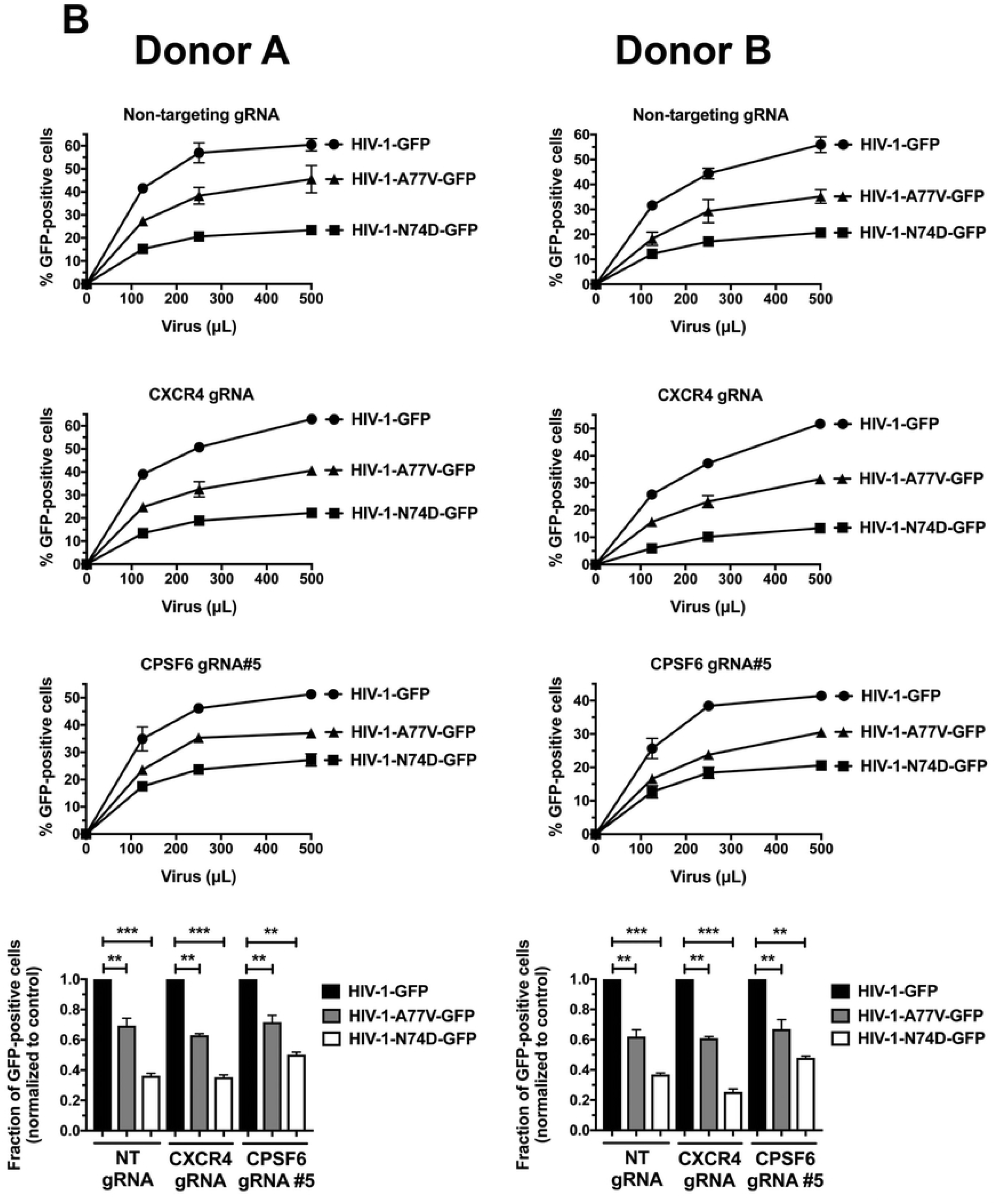
Depleted CPSF6 expression in human primary CD4^+^ T cells does not affect HIV-1 infectivity. **(A)** Human primary CD4^+^ T cells from two different donors had CPSF6 expression knocked out using the CRISPR/Cas9 system, as described in Methods. Briefly, CD4^+^ T cells were electroporated using two different guide RNAs (gRNAs) against CPSF6 (gRNA #5 and #6) together with the Cas9 protein. At 72 h post-electroporation, endogenous expression of CPSF6 in CD4^+^ T cells was analyzed by western blotting using an antibody against CPSF6. For controls, a gRNA against CXCR4 and a non-targeting gRNA were electroporated. Expression of GAPDH was used as a loading control. Similar results were obtained using two different donors, and a representative blot is shown. **(B)** Human primary CD4^+^ T cells depleted for CPSF6 expression were challenged with increasing amounts of p24-normalized HIV-1-GFP, HIV-1-N74D-GFP, or HIV-1-A77V-GFP. Infectivity was determined at 72 h post-challenge by measuring the percentage of GFP-positive cells. Experiments were repeated three times per donor, and a representative experimental result is shown. Statistical analysis was performed using an intermediate value taken from the infection curves (bottom panels). ** indicates P-value < 0.001, *** indicates P-value < 0.0005 as determined by using the unpaired t-test.

### Depletion of TRIM5α_hu_ in human primary CD4^+^ T cells rescues HIV-1-N74D infectivity

We and others have previously demonstrated that Cyp A protects the HIV-1 core from TRIM5α_hu_ restriction in human primary CD4^+^ T cells [11, 12]. Therefore, we hypothesized that TRIM5α_hu_ may decrease the infectivity of HIV-1-N74D in human primary cells. To test this hypothesis, we challenged TRIM5α_hu_-depleted human primary CD4^+^ T cells with HIV-1-N74D-GFP. crRNPs containing the anti-TRIM5α_hu_ gRNA #6 and #7 completely depleted the expression of endogenous TRIM5α_hu_ in human primary CD4^+^ T cells (Figure 4A). As shown in Figure 4B, the depletion of TRIM5α_hu_ rescued HIV-1-N74D-GFP infectivity in CD4^+^ T cells. These results suggested that TRIM5α_hu_ restricted HIV-1-N74D in human CD4^+^ T cells. Interestingly, small infectivity changes were observed for HIV-1-A77V in TRIM5α_hu_-depleted cells, suggesting that this virus is not restricted by TRIM5α_hu._

**Figure 4.**
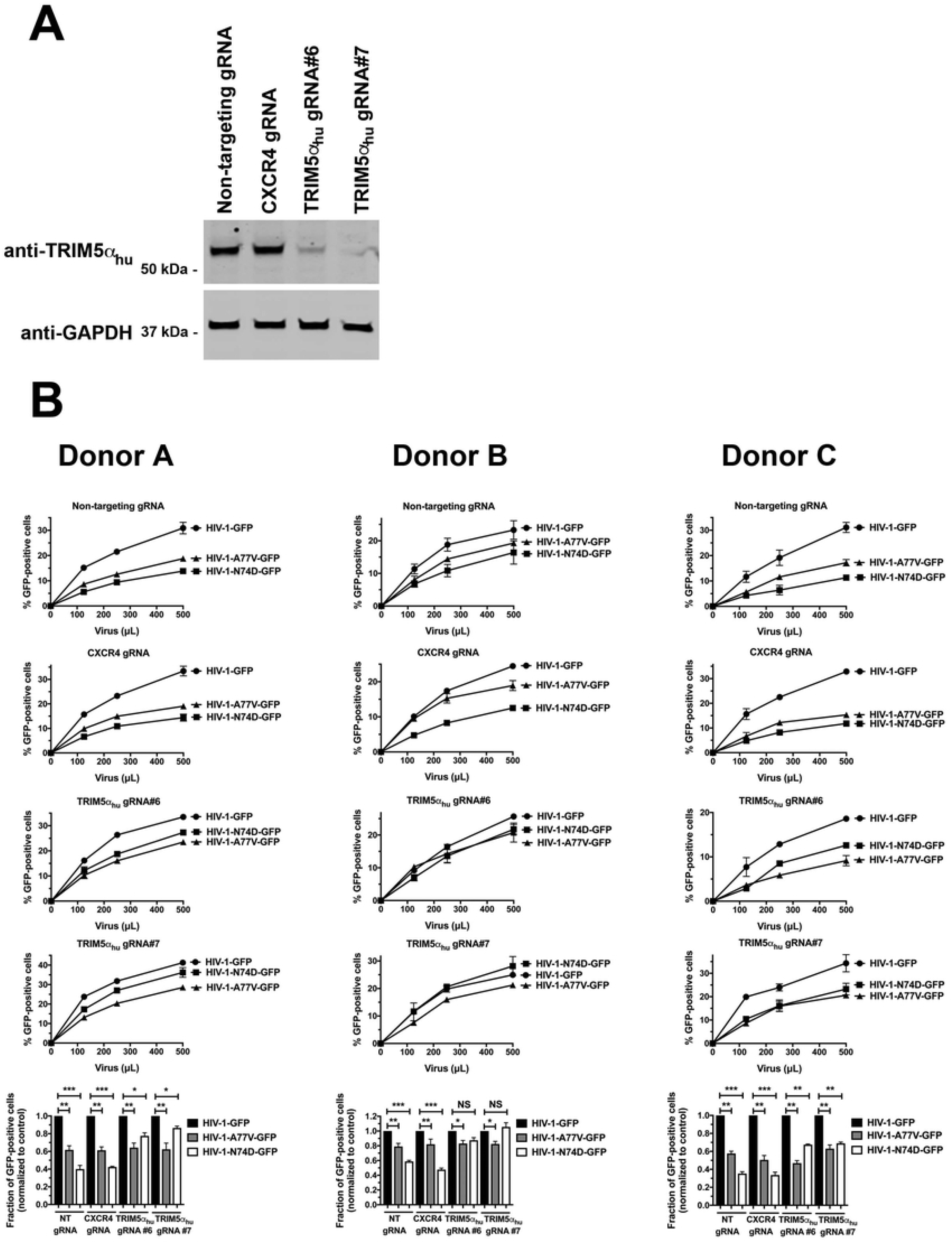
TRIM5α_hu_ depletion in human primary CD4^+^ T cells rescues HIV-1-N74D infectivity. **(A)** Human primary CD4^+^ T cells from three different donors had TRIM5α_hu_ expression knocked out using the CRISPR/Cas9 system, as described in Methods. Briefly, CD4^+^ T cells were electroporated using two different guide RNAs (gRNAs) against TRIM5α_hu_ (gRNA #6 and #7) together with the Cas9 protein. At 72 h post-electroporation, the endogenous expression of TRIM5α_hu_ in CD4^+^ T cells was analyzed by western blotting using an antibody against TRIM5α_hu_. For controls, a gRNA against CXCR4, and a non-targeting gRNA were electroporated. Expression of GAPDH was used as a loading control. Similar results were obtained using two different donors, and a representative blot is shown. **(B)** Human primary CD4^+^ T cells depleted for TRIM5α_hu_ expression were challenged with increasing amounts of p24-normalized HIV-1-GFP, HIV-1-N74D-GFP, or HIV-1-A77V-GFP. Infectivity was determined at 72 h post-challenge by measuring the percentage of GFP-positive cells. Experiments were repeated three times per donor, and a representative experimental result is shown. Statistical analysis was performed using an intermediate value taken from the infection curves (bottom panels). * indicates P-value < 0.005, ** indicates P-value < 0.001, *** indicates P-value < 0.0005, NS indicates not significant as determined by using the unpaired t-test.

### N74D-stabilized capsids bind to TRIM5α_hu_ but do not interact with Cyp A

If TRIM5α_hu_ restriction occurs after cells are infected by HIV-1-N74D, it implies that Cyp A is no longer protecting the core. To test this hypothesis, we assessed the abilities of TRIM5α_hu_ and Cyp A to bind to N74D-stabilized capsid tubes using a capsid binding assay [13]. As shown in Figure 5, TRIM5α_hu_ bound with increased affinity to stabilized N74D capsid tubes than to wild-type tubes. Interestingly, TRIM5α_hu_ bound to A77V-stabilized tubes and wild-type tubes in a similar manner. Results from these ancillary experiments support the idea that the infectivity defect observed for HIV-1-N74D is due to an increase in TRIM5α_hu_ binding to N74D capsids compared with that to wild-type capsids. We have previously shown that when Cyp A was not expressed in primary CD4^+^ T cells, TRIM5α_hu_ binding to capsid increased [11]; therefore, we tested the ability of Cyp A to bind to N74D-stabilized capsid tubes. As shown in Figure 5, Cyp A did not bind to N74D-stabilized capsid tubes, although it bound to wild-type capsid tubes. These results showed that N74D capsids were not protected by Cyp A, leading to TRIM5α_hu_ binding and restriction. Interestingly, Cyp A did not bind to A77V-stabilized tubes.

**Figure 5.**
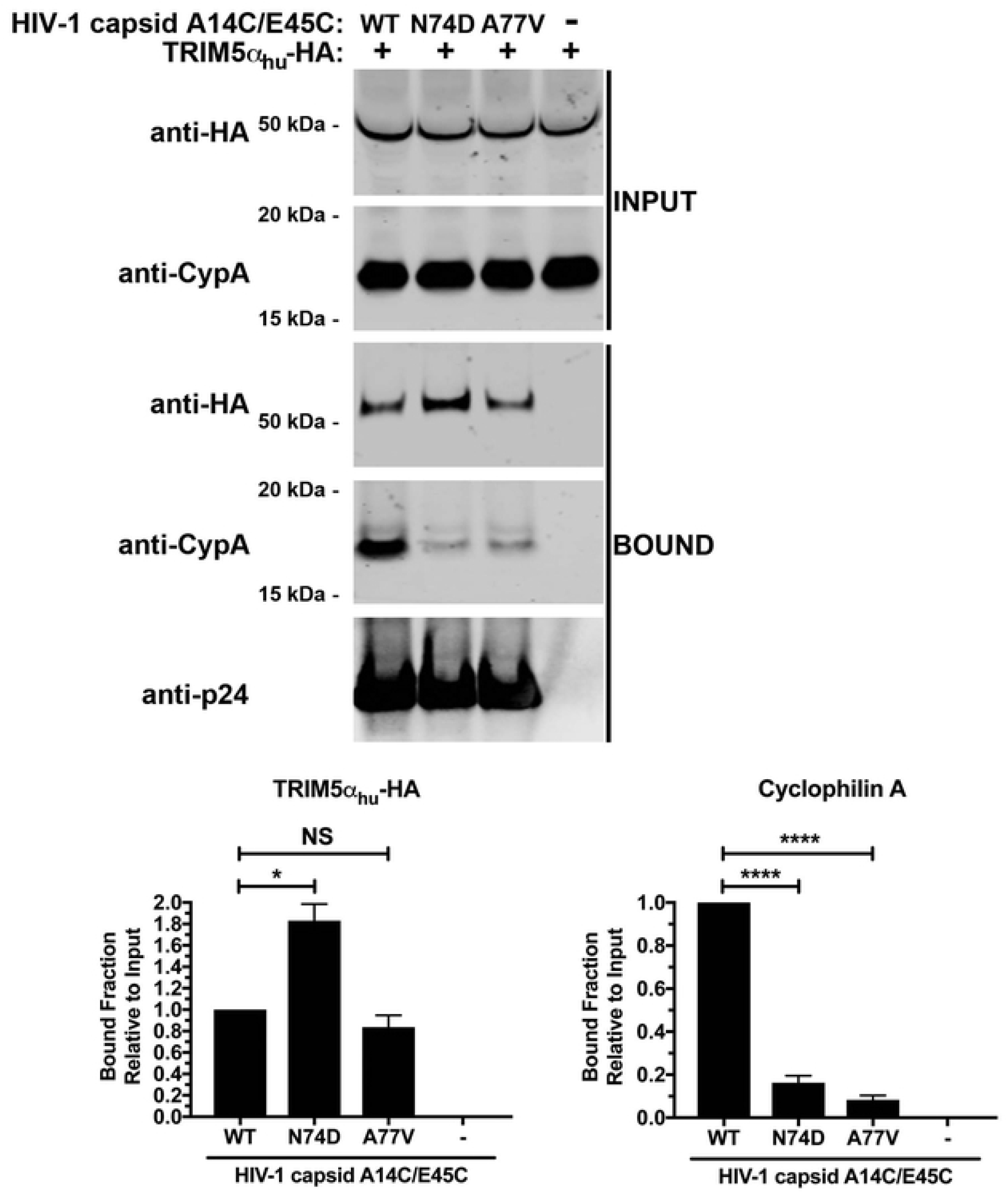
N74D-stabilized capsids bind to TRIM5α_hu_ but do not interact with Cyp. **A**. Human 293T cells were transfected with plasmids expressing TRIM5α_hu_- hemagglutinin (HA). Post-transfection (24 h), cells (INPUT) were lysed in capsid binding buffer (CBB) as described in Methods. Cell extracts containing TRIM5α_hu_-HA were then mixed with 10 μl of either stabilized wild-type, N74D, or A77V capsid tubes (5 mg/ml). Mixtures were incubated for 1 h at room temperature. Stabilized HIV-1 capsid tubes were collected by centrifugation and washed twice using CBB. Pellets were resuspended in 1× Laemmli buffer (BOUND). INPUT and BOUND fractions were then analyzed by western blotting using anti-HA, anti-Cyp A, and anti-p24 antibodies. Experiments were repeated three times, and a representative experimental result is shown. The BOUND fraction relative to the INPUT fraction for three independent experiments (with standard deviation) is shown. * indicates a p-value < 0.005, **** indicates a p-value < 0.0001, and NS indicates no significant difference as determined by unpaired t-tests.

## DISCUSSION

Here we have shown that the infectivity defect observed for HIV-1-N74D viruses in CD4^+^ T cells is due to its inability to interact with Cyp A, which exposes the viral core to TRIM5α_hu_ binding and restriction. These results concur with the idea that Cyp A plays an important role in the protection of the HIV-1 core in the early stages of infection [11, 12], and indicate once again that Cyp A may be essential in the early stages of HIV-1 infection to ensure protection of the core from restriction factors or cellular conditions that may affect infection. These results are in contrast to the current notion that the infectivity defect of N74D is due to its loss of CPSF6 binding. Although capsids bearing the N74D change do not interact with CPSF6, the infectivity defect that HIV-1-N74D viruses exhibit is due to a decrease in Cyp A binding with a concomitant gain of TRIM5α_hu_ binding, which restricts infection.

While an intact HIV-1 core has more than ∼1200 binding sites for Cyp A, the actual number of sites occupied by Cyp A during an infection is not known; however, it is reasonable to thing that binding of one or two Cyp A molecules per hexamer would be sufficient to prevent the binding of restriction factors such as TRIM5α_hu_ by steric hindrance. So theoretically, only two Cyp A molecules per hexamer would be needed to ensure that infection is productive. Interestingly, residue N74 in the capsid structure is distantly located from the Cyp A binding loop, suggesting that an overall structural shift may be occurring in order to prevent Cyp A binding. An alternative explanation is that the N74D mutation may affect core breathing, and consequently, inhibit the binding of Cyp A to the core [14]. One of the implications of this study is that interactions between Cyp A and capsid mutants should be considered when trying to understand HIV-1 infectivity defects involving human primary cells.

The infectivity defect of HIV-1-A77V viruses was not very pronounced when compared to HIV-1-N74D viruses. In addition, depletion of TRIM5α_hu_ did not rescue the infectivity defect of HIV-1-A77V. In agreement, A77V stabilized capsid tubes did not showed an increase in binding to TRIM5α_hu_, but lost binding to Cyp A. One possibility is that HIV-1-A77V viruses are defective for a different reason, which agrees with the experiments showing that HIV-1-A77V viruses can replicate in CD4+ T cells when compared to HIV-1-N74D [15].

While this work was ongoing, TRIM34 has also been shown to be important for decreased HIV-1-N74D infectivity in primary CD4^+^ T cells [16]. Taken together, these results suggest that TRIM5α_hu_ may be working together with TRIM34 to reduce HIV-1-N74D infectivity. We demonstrated previously that TRIM5α proteins can form higher-order, self-associating complexes which are essential for TRIM5α-based restriction of retroviruses [17, 18]. In addition, these studies showed that TRIM5α proteins can also form higher-order complexes with TRIM orthologs such as TRIM34 and TRIM6 [19]. It is possible that TRIM5α_hu_ forms higher-order complexes with TRIM34 in order to bind and restrict HIV-1-N74D viruses, as we have previously suggested [20].

These results highlight the importance of HIV-1 core-Cyp A interactions during productive HIV-1 infection and indicate that Cyp A is an essential cofactor for HIV-1 replication in human primary CD4^+^ T cells.

## MATERIALS AND METHODS

### Infection using HIV-1-GFP reporter viruses

Recombinant HIV-1 strains (e.g., HIV-1-N74D and HIV-1-A77V) expressing GFP, and pseudotyped with VSV-G, were prepared as previously described [21]. All viruses were titered according to p24 levels and infectivity. Viral challenges were performed in 24-well plates by infecting 50,000 cells (PBMCs or CD4^+^ T cells) per well. Infectivity was determined by measuring the percentage of GFP-positive cells using flow cytometry (BD FACSCelesta, San Jose, CA, USA).

### Capsid expression and purification

The pET-11a vector was used to express HIV-1 capsid proteins containing the A14C and E45C mutations. Point mutations N74D and A77V were introduced using the QuikChange II site-directed mutagenesis kit (Stratagene) according to the manufacturer’s instructions. All proteins were expressed in *Escherichia coli* one-shot BL21star (DE3) cells (Invitrogen, Carlsbad, CA, USA), as previously described [13]. Briefly, cells were inoculated in Luria-Bertani medium and cultured at 30°C until mid-log phase (Absorbance at 600 nm, 0.6–0.8). Protein expression was induced with 1 mM isopropyl-β-d-thiogalactopyranoside overnight at 18°C. Cells were harvested by centrifugation at 5,000 × g for 10 min at 4°C, and pellets were stored at −80°C until purification. Purification of capsids was carried out as follows. Pellets from two-liter cultures were lysed by sonication (Qsonica microtip: 4420; A = 45; 2 min; 2 sec on; 2 sec off for 12 cycles), in 40 ml of lysis buffer (50 mM Tris pH = 8, 50 mM NaCl, 100 mM β-mercaptoethanol, and Complete ethylenediaminetetraacetic acid (EDTA)-free protease inhibitor tablets). Cell debris was removed by centrifugation at 40,000 × g for 20 min at 4°C. Proteins from the supernatant were precipitated by incubation with one-third the volume of saturated ammonium sulfate containing 100 mM β-mercaptoethanol for 20 min at 4°C, and centrifugation at 8,000 × g for 20 min at 4°C. Precipitated proteins were resuspended in 30 ml of buffer A (25 mM 2-(N-morpholino) ethanesulfonic acid (MES), pH 6.5, and 100 mM β-mercaptoethanol) and sonicated 2–3 times (Qsonica microtip: 4420; A = 45; 2 min; 1 sec on; 2 sec off). The protein sample was dialyzed three times in buffer A (2 h, overnight, and 2 h), sonicated, diluted in 500 ml of buffer A, and then separated sequentially on a 5-ml HiTrap Q HP column followed by a 5-ml HiTrap SP FF column (GE Healthcare), which were both pre-equilibrated with buffer A. Capsid proteins were eluted from the HiTrap SP FF column using a linear gradient of concentrations ranging from 0–2 M NaCl. The eluted fractions that had the highest protein levels were selected based on absorbance at 280 nm. Pooled fractions were dialyzed three times (2 h, overnight, and 2 h) in storage buffer (25 mM MES, 2 M NaCl, 20 mM β-mercaptoethanol). Samples were concentrated to 20 mg/ml using Centriprep Centrifugal Filter Units and stored at –80°C.

### Assembly of stabilized HIV-1 capsid tubes

One milliliter of monomeric capsid (5 mg/ml) was dialyzed in SnakeSkin dialysis tubing (10K MWCO, Thermo Scientific, Waltham, MA, USA) using a buffer that was high in salt and contained a reducing agent (Buffer 1: 50 mM Tris, pH 8, 1 M NaCl, 100 mM β-mercaptoethanol) at 4°C for 8 h. The protein was then dialyzed using the same buffer without the reducing agent β-mercaptoethanol (Buffer 2: 50 mM Tris, pH 8, 1 M NaCl) at 4°C for 8 h. The absence of β-mercaptoethanol in the second dialysis allowed the formation of disulfide bonds between Cysteine 14 and the 45 inter-capsid monomers in the hexamer. Finally, the protein was dialyzed using Buffer 3 (20 mM Tris, pH 8, 40 mM NaCl) at 4°C for 8 h. Assembled complexes were kept at 4°C for up to 1 month.

### Capsid binding assay protocol

Human HEK293T cells were transfected for 24 h with a plasmid expressing the protein of interest (TRIM5α_hu_). The culture media was completely removed and cells were scraped off the plate and lysed in 300 μl of capsid binding buffer (CBB: 10 mM Tris, pH 8, 1.5 mM MgCl_2_, 10 mM KCl). Cells were rotated for 15 min at 4°C and then centrifuged to remove cellular debris (21,000 *×* g, 15 min, 4°C). Cell lysates were incubated with stabilized HIV-1 capsid tubes for 1 h at 25°C. The stabilized HIV-1 capsid tubes were then centrifuged at 21,000 × g for 2 min. Pellets were washed 2–3 times by resuspension and centrifugation in CBB or phosphate-buffered saline (PBS). Pellets were resuspended in 1× Laemmli buffer and analyzed by western blotting using an anti-p24 antibody and other appropriate antibodies.

### Preparation of PBMCs and CD4^+^ T cells

PBMCs from healthy-donor whole blood were isolated by density gradient centrifugation using Ficoll-Paque Plus (GE Health Care, Chicago, IL, USA). Whole blood (40 ml) was centrifuged at 300 × g for 10 min, and the plasma layer was removed and replaced with Hank’s Balanced Salt solution (HBSS; Sigma Aldrich, St. Louis, MO, USA). The blood sample was then diluted 1:2 with HBSS, and 20 ml of the diluted sample was layered on top of 20 ml Ficoll-Paque Plus and centrifuged at 300 × g for 30 min. The resulting buffy coat layer was collected, washed twice with HBSS, and resuspended in Roswell Park Memorial Institute (RPMI) medium containing 10% (vol/vol) fetal bovine serum (FBS) and 1% (vol/vol) penicillin-streptomycin, and activated with IL-2 (100 U/ml) (Human IL-2; Cell Signaling Technology, #8907SF) and phytohemagglutinin (1 µg/ml) for 3 days. CD4^+^ T cells were obtained via negative selection from PBMCs using a human CD4^+^ T-cell isolation kit (MACS Miltenyi Biotec, #130-096-533, Bergisch Gladbach, Germany). PBMCs (1 × 10^7^ total cells) were resuspended in 40 µl of CD4^+^ T-cell isolation buffer (PBS, pH 7.2, 0.5% bovine serum albumin (BSA), and 2 mM EDTA). CD4^+^ T-cell biotin-antibody cocktail (10 µl) was then added to the PBMCs and incubated at 4°C for 5 min. CD4^+^ T-cell isolation buffer (30 µl) and CD4^+^ T-cell microbead cocktail (20 µl) were then added and further incubated for 10 min at 4°C. Depending on the number of PBMCs isolated, either an LS or MS column attached to a Magnetic Activated Cell Sorting Separator was prewashed using 3 ml or 6 ml of ice-cold CD4^+^ T-cell isolation buffer, respectively. The PBMC suspension was added to the column and the flow-through was collected in a 15-ml tube. The LS or MS column was then washed (3 ml or 6 ml, respectively, with ice-cold CD4^+^ T-cell isolation buffer), and the flow-through was collected. The newly isolated CD4^+^ T cells were then centrifuged at 800 × g for 5 min and resuspended in RPMI medium supplemented with IL-2 (100 U/ml).

### CRISPR-Cas9 knockouts in primary CD4^+^ T cells

Detailed protocols for the production of CRISPR-Cas9 ribonucleoprotein complexes (crRNPs) and primary CD4^+^ T-cell editing have been previously published [22, 23]. Briefly, lyophilized CRISPR RNA (crRNA) and trans-activating crispr RNA (tracrRNA; Dharmacon, Lafayette, CO, USA) were each resuspended at a concentration of 160 µM in 10 mM Tris-HCl (pH 7.4), and 150 mM KCl. Five microliters of 160 µM crRNA was then mixed with 5 µl of 160 µM tracrRNA and incubated for 30 min at 37°C. The gRNA:tracrRNA complexes were then mixed gently with 10 µl of 40 µM purified Cas9-NLS protein (UC-Berkeley Macrolab) to form crRNPs. Complexes were aliquoted and frozen in 0.2-ml PCR tubes (USA Scientific, Ocala, FL, USA) at – 80°C until further use. crRNA guide sequences used in this study were a combination of sequences derived from the Dharmacon predesigned Edit-R library for gene knockouts, and custom-ordered sequences as indicated.

PBMCs were isolated by density gradient centrifugation using Ficoll-Paque Plus (GE Health Care, #17-1440-02). PBMCs were washed thrice with 1× PBS to remove platelets and resuspended at a final concentration of 5 × 10^8^ cells/ml in 1× PBS, 0.5% BSA, and 2 mM EDTA. Bulk CD4^+^ T cells were subsequently isolated from PBMCs by magnetic negative selection using an EasySep Human CD4+ T Cell Isolation Kit (STEMCELL, per manufacturer’s instructions). Isolated CD4^+^ T cells were suspended in complete RPMI medium, consisting of RPMI-1640 (Sigma Aldrich) supplemented with 5 mM 4-(2-hydroxyethyl)-1-piperazineethanesulfonic acid (Corning, Corning, NY, USA), 50 μg/ml penicillin/streptomycin (Corning), 5 mM sodium pyruvate (Corning), and 10% FBS (Gibco). Media was supplemented with 20 IU/ml IL-2 (Miltenyi) immediately before use. For activation, bulk CD4^+^ T cells were immediately plated on anti-CD3-coated plates [coated for 12 hours at 4°C with 20 µg/ml anti-CD3 antibody (UCHT1, Tonbo Biosciences)] in the presence of 5 µg/ml soluble anti-CD28 antibody (CD28.2, Tonbo Biosciences). Cells were stimulated for 72 h at 37°C in a 5%-CO_2_ atmosphere prior to electroporation. After stimulation, cell purity and activation were verified by CD4/CD25 immunostaining and flow cytometry as previously described [22].

After three days of stimulation, cells were resuspended and counted. Each electroporation reaction consisted of between 5 × 10^5^ and 1 × 10^6^ T cells, 3.5 µl RNPs, and 20 µl of electroporation buffer. crRNPs were thawed to room temperature. Immediately prior to electroporation, cells were centrifuged at 400 × g for 5 minutes, the supernatant was removed by aspiration, and the pellet was resuspended in 20 µl of room temperature P3 electroporation buffer (Lonza, Basel, Switzerland) per reaction. Cell suspensions (20 µl) were then gently mixed with each RNP and aliquoted into a 96-well electroporation cuvette for nucleofection with the 4D 96-well shuttle unit (Lonza) using pulse code EH-115. Immediately after electroporation, 100 µl of prewarmed media without IL-2 was added to each well and cells were allowed to rest for 30 min in a cell-culture incubator at 37°C. Cells were subsequently moved to 96-well flat-bottomed culture plates prefilled with 100 µl warm complete media with IL-2 at 40 U/ml (for a final concentration of 20 U/ml) and anti-CD3/anti-CD2/anti-CD28 beads (T cell Activation and Stimulation Kit, Miltenyi) at a 1:1 bead:cell ratio. Cells were cultured at 37°C in a 5%-CO_2_ atmosphere, dark, humidified cell-culture incubator for four days to allow for gene knockout and protein clearance, with additional media added on day 2. To check knockout efficiency, 50 µl of mixed culture was transferred to a centrifuge tube. Cells were pelleted, the supernatant removed, and the pellets were resuspended in 100 µl 2.5× Laemmli Sample Buffer. Protein lysates were heated to 98°C for 20 min before storage at –20°C until assessment by western blotting.

RNA guides:

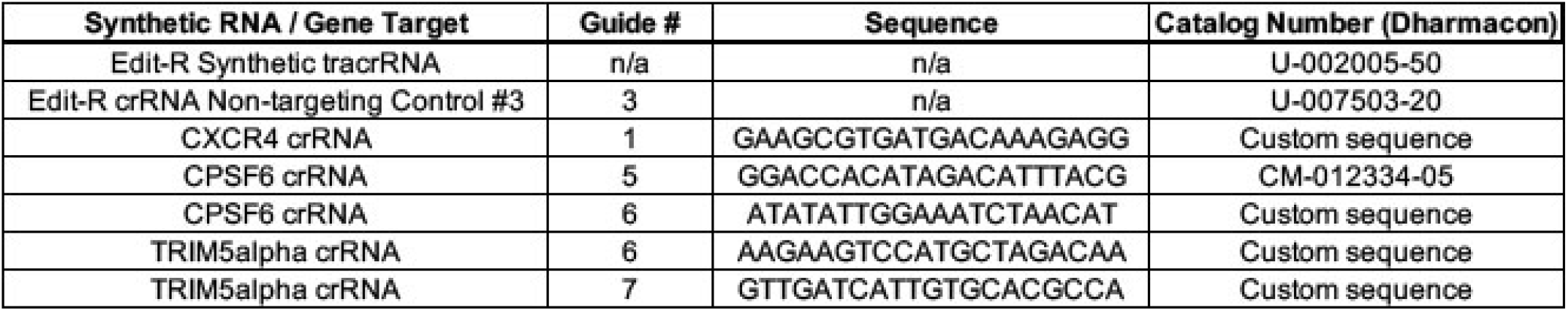

## QUANTIFICATION AND STATISTICAL ANALYSES

Statistical analyses were performed using unpaired t-tests. Sample numbers, number of replicates, and *p* values are indicated in corresponding figure legends. Quantification of western blot band intensities was performed using ImageJ. For all experiments, means and standard deviations were calculated using GraphPad Prism 7.0c.

## ACKNOWLDGEMENTS

We thank the NIH AIDS repository for reagents. A.S. and F.D.-G. are supported by an AI087390 grant to F.D.-G.

